# The evolution of allostery in a protein family

**DOI:** 10.1101/2025.06.20.660748

**Authors:** Aina Martí-Aranda, Ben Lehner

**Affiliations:** Wellcome Sanger Institute, Cambridge, UK; Centre for Genomic Regulation (CRG), Barcelona Institute for Science and Technology (BIST), Barcelona, Spain; Universitat Pompeu Fabra (UPF), Barcelona, Spain; Institució Catalana de Recerca i Estudis Avançats (ICREA), Barcelona, Spain

## Abstract

Allosteric interactions in proteins are key to biological regulation and the efficacy of many drugs. The extent to which allostery is conserved in evolutionarily-related proteins is unknown, with implications for predicting, engineering and therapeutically targeting allostery. Here we directly address this question by constructing seven comprehensive allosteric maps for five homologous human proteins. The comparative maps reveal a modular allosteric architecture with conserved distant-dependent allosteric decay across the protein core connecting to protein-specific allosteric extensions to surfaces. These allosteric augmentations use both structurally-conserved residues and homolog-specific domain extensions. Our data provide the first comparative multidimensional protein energy landscapes and suggest that allostery evolves via the gain-and-loss of peripheral extensions to a conserved allosteric core.

## Introduction

The transmission of information spatially from one site to another in a protein is termed allostery^1^. Allostery is central to biological regulation, allowing protein activities to be controlled by covalent modifications, the binding of other molecules, and mutations outside of active sites. As a consequence, Monod infamously termed allostery ‘the second secret of life’^2,3^. Many disease-causing mutations, including cancer driver mutations, are pathogenic because of their allosteric effects. Reciprocally, many effective therapeutics target allosteric sites^4^. Allosteric drugs can have higher specificity than orthosteric drugs targeting highly-conserved active sites. Moreover they also allow activation of proteins and finer control, modulating the response to endogenous regulation rather than blocking catalytic centres^4^

It is often stated that allosteric sites are less conserved than active sites^5,6^. Whilst there is evidence that this is true at the sequence level^7,8^, the structural conservation of allosteric sites is less clear^6^, and the functional conservation of allostery in proteins is, to our knowledge, largely unknown. The reason for this is simple: methods to comprehensively quantify allosteric communication in proteins have only recently been developed and no comprehensive allosteric maps have, to date, been produced for any homologous proteins. As such, we do not know the extent to which the same positions in evolutionarily-related proteins have allosteric effects when perturbed.

At one extreme it could be that allosteric coupling is highly conserved in proteins with the same fold. At the other extreme, allostery may be fast evolving, with each member of a protein family having different allosteric sites. An alternative possibility is that allostery is conserved in a subset of sites but fast evolving in others. Based on testing individual mutations, both conservation and divergence of allosteric effects has been reported^9^.

A second interesting question about allostery is how the addition of extra residues to a protein couples to the existing allosteric network. Domains are considered evolutionary modules of proteins that can fold and function independently. However, individual protein domains quite frequently have additional secondary structure elements^10,11^ and each protein typically consists of multiple domains as well as more dynamic intrinsically disordered regions. In at least some cases, these extensions can have important allosteric effects^11^.

The extent to which allostery is conserved in evolutionarily-related proteins has important implications for predicting, engineering, and therapeutically targeting allosteric sites. For example, if the accessible allosteric sites differ among members of a protein family this should allow therapeutics binding these sites to specifically inhibit or activate one member of the family, even if they physically bind to multiple homologous proteins.

Here we directly test the conservation of allostery in homologous proteins by generating seven comprehensive allosteric maps for five homologous human protein domains. As a model protein family we use PDZ domains (Fig. 1a). The domain is named for three proteins—postsynaptic density protein (PSD-95), Drosophila discs large tumor suppressor (DlgA), and zonula occludens-1 protein (ZO-1)^12^. PDZ domains are the largest family of human protein-protein interaction domains^13^ and so provide an ideal model system for investigating energetic and allosteric conservation in homologous proteins. More than 270 PDZ domains in 155 different proteins participate in a wide range of cellular processes, including signal transduction, cell polarity, cell adhesion, and neuronal synaptic transmission, often functioning as scaffolds that mediate protein-protein assembly^13^. Many of these domains are considered high-value therapeutic targets, but very few have been successfully targeted^14,15^. Despite low sequence identity, PDZ domains share a canonical fold composed of five to six β-strands and two or three α-helices. PDZ domains bind diverse, predominantly C-terminal peptide ligands, often classified into three classes by the identity of ligand positions, based on ligand residues at positions 0 and -2 (position 0 is the C-terminal residue; position -1 is the residue before)^16,17^. Ligand recognition occurs in a pocket composed of the β2 strand, the α2 helix and the carboxylate-binding β1-β2 loop^17,18^ (Fig. 1b).

**Figure 1:**
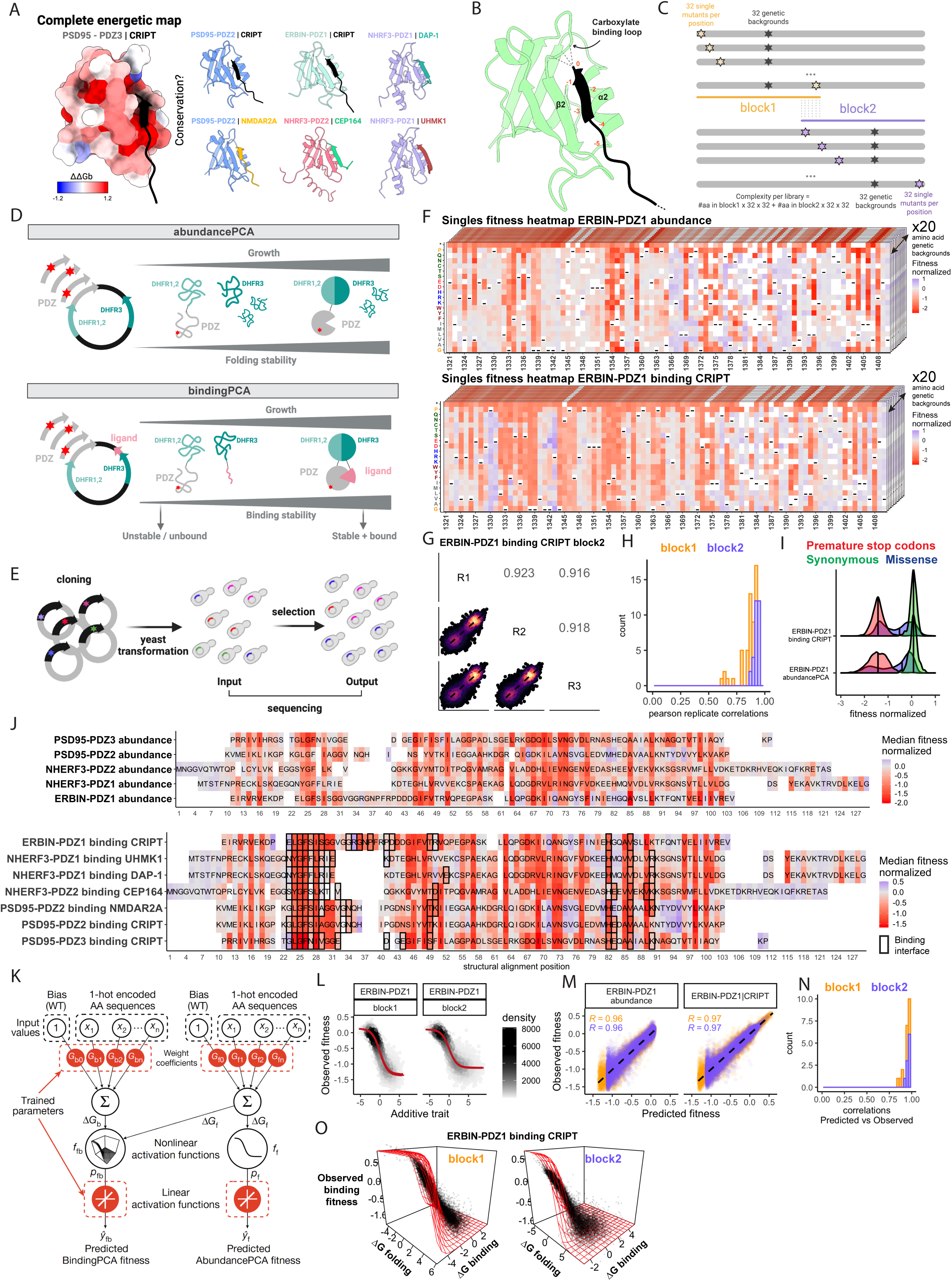
Seven complete energetic and allosteric landscapes for PDZ domains. **a.** Energetic surface map for PSD95-PDZ3 binding CRIPT ^19^ alongside six homologous interactions. Structures are AlphaFold3 ^44^ predictions. **b.** Canonical fold and binding of the PDZ domain family (PSD95-PDZ2 binding CRIPT). Ligand residues numbered in orange. **c.** Library design. For each PDZ domain, two overlapping library blocks, incorporating NNK codons at every position and an NNK codon in a fixed position. Schematics were created using BioRender.com. **d.** AbundancePCA (aPCA) and BindingPCA (bPCA) selections. Schematics were created using BioRender.com. **e.** Deep mutational scanning experimental workflow. Schematics were created using BioRender.com. **f.** Heat maps for fitness effects of single amino acid substitutions for ERBIN-PDZ1 from aPCA (top) and bPCA (bottom). Dashes indicate wild-type residues. Residue numbering from UniProt. Stacked heat maps represent data across 20 genetic backgrounds. **g.** Example of triplicate fitness Pearson correlations for ERBIN-PDZ1 binding CRIPT (library block 2). **h.** Distribution of Pearson correlation coefficients for triplicate fitness measurements for all libraries. **i.** Example of fitness density distributions for libraries for ERBIN-PDZ1. Fitness measurements shown after normalization between blocks and libraries. **j.** Median of the normalised fitness values mapped onto PDZ structural alignment. Residue numbering corresponds to alignment positions. **k.** Neural network architecture used to fit thermodynamic models to the PCA data (bottom, target and output data), thereby inferring the causal changes in free energy of folding and binding associated with single amino acid substitutions (top, input values). **l.** Example of non-linear relationships (global epistasis) between observed aPCA fitness and changes in free energy of folding for EBRIN-PDZ1. **m**. Examples of performance of models fit to PCA data for EBRIN-PDZ1 libraries. **n.** Distribution of Pearson correlation coefficients for MoCHI observed vs predicted fitness measurements. **o**. Example of non-linear relationships (global epistasis) between observed BindingPCA fitness and both changes in free energy of folding and binding for EBRIN-PDZ1 binding CRIPT.

The seven comprehensive energetic and allosteric maps reported here allow comparative analysis of homologous binding interfaces and reveal a modular evolutionary architecture of allostery, with a conserved allosteric core connected to protein-specific allosteric extensions. We propose a model whereby allostery evolves via the gain-and-loss of peripheral extensions to a conserved allosteric core.

## Results

### Generating seven complete allosteric and energetic maps

To test the conservation of the energetic effects of mutations and allostery in homologous proteins we quantified changes in folding energy and binding energy for all mutations in five different human PDZ domains and a total of seven different protein interactions (Fig. 1a). The five human PDZ domains - PSD95-PDZ3, PSD95-PDZ2, NHERF3-PDZ1, NHERF3-PDZ2, and ERBIN-PDZ1 (Fig. 1a) - have 29.6% mean sequence identity (range from 23.5% to 38.1%) and mean backbone RMSD of 2.61Å in pairwise structural alignments (range 1.85Å to 5.65Å). Our experimental design has a three tiered structure, quantifying the energetic effects of mutations on the same PDZ domain binding to different ligands (PSD95-PDZ2 binding peptides from CRIPT and NMDAR2A; NHERF3-PDZ1 binding peptides from UHMK1 and DAP-1), different PDZ domains binding to the same ligand (PSD95-PDZ3, PSD95-PDZ2, NHERF3-PDZ1, and ERBIN-PDZ1 binding CRIPT), and different PDZ domains binding different ligands. All ligands are class 1 ligands, except CEP164 which is a class 2 ligand^17^ (see methods for the design strategy). The data for six interactions were generated for this study, with the data for PSD95-PDZ3 previously reported in Faure et al. 2022^19^.

For each PDZ domain we generated two overlapping libraries containing both single and double amino acid (aa) substitutions using NNK (N=[A,G,T,C], K=[G,T]) codons to encode all 20 amino acids at each position in combination with a second NNK codon to encode all 20 aa at a second fixed site (Fig. 1c and Methods). The effect of each aa substitution at each position is thus tested alone and in combination with 19 additional substitutions. As previously described, quantifying how mutations interact in double mutants allows us to infer the underlying additive causal free energy changes for each variant^19–23^.

To quantify the cellular abundance and peptide binding of each variant we used two extensively validated pooled protein-fragment complementation assays (PCA)^19,24,25^. In the first assay, AbundancePCA (aPCA), variants are expressed in yeast cells fused to a short fragment of the enzyme dihydrofolate reductase (DHFR), with the concentration of the fusion protein controlling the cellular growth rate^19,22,24^. In the second assay, BindingPCA (bPCA), the PDZ variants are fused to one fragment of DHFR and the peptide ligand is expressed fused to the remaining DHFR sequence such that the cellular growth rate is controlled by the concentration of the protein-peptide complex^19,24^ (Fig. 1d). Mutational effects on protein abundance and binding are quantified using sequencing to quantify the change in frequency of each variant during selection (Fig. 1e).

In total, the data analysed here derive from 22 different pooled selections with three replicate abundance measurements for a total of 172,200 amino acid (424,512 nucleotide) genotypes and triplicate binding measurements for 248,200 amino acid (619,072 nucleotide) genotypes. After selections and quality filtering (see Methods) we obtained reliable abundance and binding measurements for a total of 153,562 and 241,200 protein variants, respectively (Fig 1f, Extended Data Fig. 1c and Supplementary Table 4).

The measurements of abundance and binding were highly reproducible, with median Pearson correlation coefficient, *r* = 0.88 and 0.92 for abundance and binding, respectively (Fig. 1f,g, and Extended Data Fig. 1a), excellent block measurements overlap (Fig1c, and Extended Data Fig. 1b), and excellent separation of synonymous and stop variants (Fig. 1h and Extended Data Fig. 1a).

Plotting binding against abundance shows, as expected^19,22^, that many changes in binding are caused by reduced protein abundance. However, a large number of mutations in each protein have larger effects on binding than can be explained by changes in abundance, particularly for mutations in the binding interface (Extended Data Fig. 1d). Quantifying the median change in abundance and binding for mutations at each position suggests conservation of mutational effects across the seven interactions (Fig. 1j, Extended Data Fig. 1c).

### From molecular phenotypes to free energy changes

We used MoCHI^20^ to fit a three-state thermodynamic model to the data for each protein to infer the underlying causal changes in folding and binding energies for each mutation. The model accounts for the non-linear relationships between the Gibbs free energy of folding (ΔG_f_) and binding (ΔG_b_) and the observed molecular phenotypes, while assuming that the energetic effects of mutations (ΔΔG_f_ and ΔΔG_b_) combine additively in double mutants. The additive energy models are highly predictive with a median *r =* 0.96 between observed and predicted fitness effects across all experiments, evaluated by ten-fold cross-validation (Fig. 1l,o, 2a, Extended Data Fig. 2a-c). The free energy measurements also agree with independent measurements made using different experiments^26^ (*r* = 0.89 for n=171 ΔΔG_b_ measurements for PSD95-PDZ2 binding NMDAR2A and *r* = 0.76 for n=168 ΔΔG_b_ measurements for PSD95-PDZ3 binding CRIPT) (Fig. 2b).

**Figure 2:**
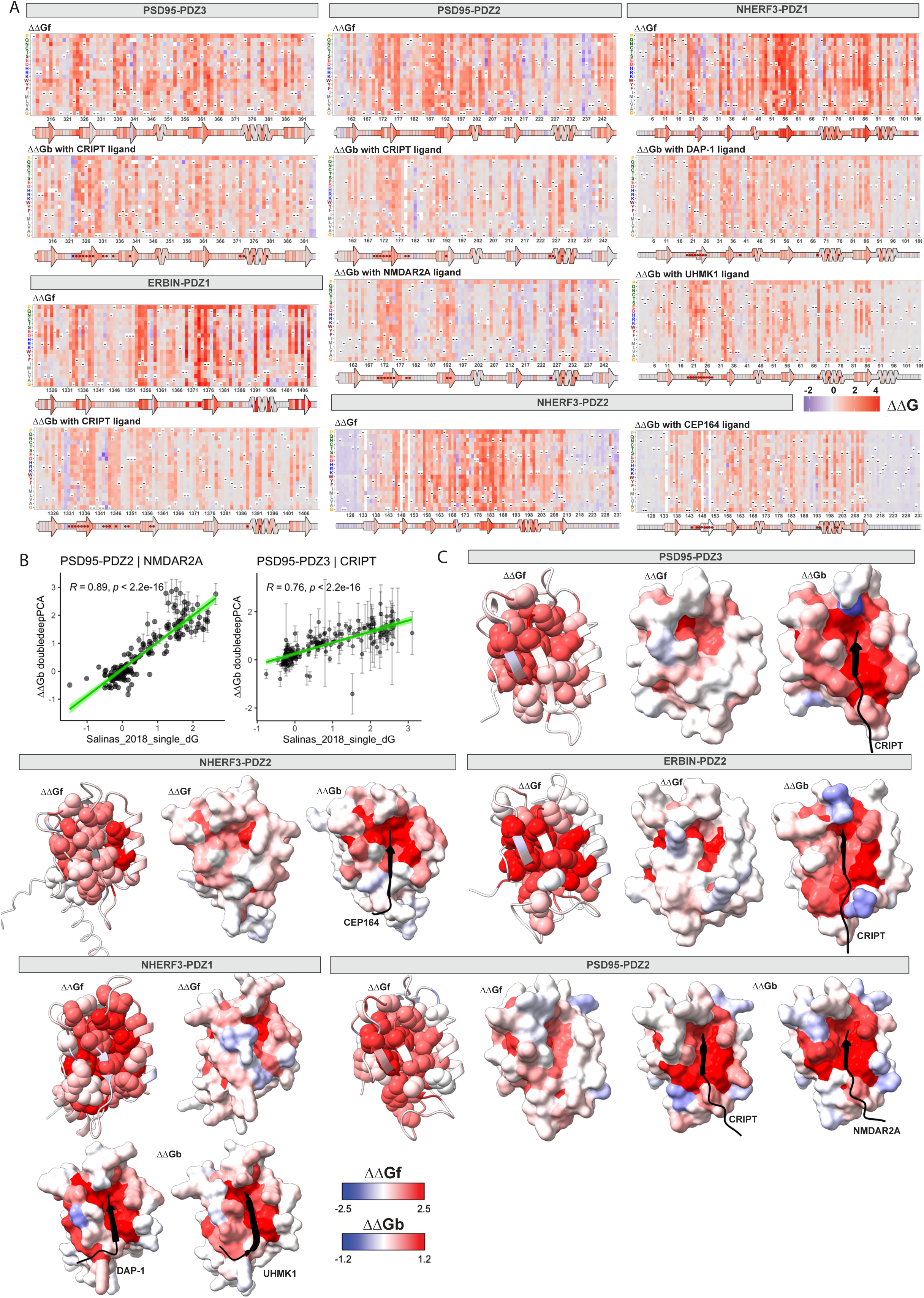
Comprehensive PDZ domain energy landscapes. **a.** Free energy heat maps for all PDZ domain interactions. Dashes indicate wild-type residues. Residue numbering follows UniProt protein sequence annotation. **b.** Correlations with independent energy measurements from Salinas and Ranganathan 2018^26^. BindingPCA free energies are from a single model; error bars indicate 95% confidence intervals from a Monte Carlo simulation approach (n = 10 experiments). Pearson’s R is shown together with a linear model fit with 95% confidence intervals shown with green shading. **c.** PDZ domain structures colored by median free energy changes (ΔΔG) at each residue position. For each domain, the first structure highlights median ΔΔG_f_ values in the core residues. The second and third structures are oriented towards the ligand-binding interface, displaying surface coloring based on ΔΔG_f_ and ΔΔG_b_, respectively.

The final dataset quantifies 21,802 free energy changes - 9,064 changes in fold stability (ΔΔG_f_) for five PDZ domains and 12,738 changes in binding energy (ΔΔG_b_) across seven interactions (Supplementary Table 5). To our knowledge, this dataset represents the first large-scale measurement of free energy changes for the interactions of homologous proteins (Fig. 2a).

Mapping the median change in free energy onto the molecular structures of the PDZ domains reveals, as expected, that mutations causing large changes in fold stability (ΔΔG_f_) are enriched in the buried protein cores whereas mutations in surface-exposed residues produce comparatively milder effects on folding (Fig. 2c, Extended Data Fig. 2d). Also as expected, mutations causing large changes in binding energy (ΔΔG_b_) are highly enriched in the binding interfaces (Fig. 2c, Extended Data Fig. 2e).

### Binding interface energy landscapes for seven interactions

The seven comprehensive mutational effect binding energy matrices provide a unique opportunity for a comparative analysis of the binding interfaces of homologous proteins. It has previously been reported that mutations in only a subset of the structural contacts in a protein interaction interface strongly affect the binding energy^22,27,28^. These sites are referred to as binding interface ‘hotspots’^27,29^.

The data for the seven PDZ domain interactions is consistent with the hotspot concept, with a bimodal distribution of ΔΔG_b_ values for mutations in each interface (defined as those residues making direct contacts with the ligand, Fig. 3a) and mutations in only a subset of these structural contacts strongly affect the binding energy (Fig. 3b).

**Figure 3:**
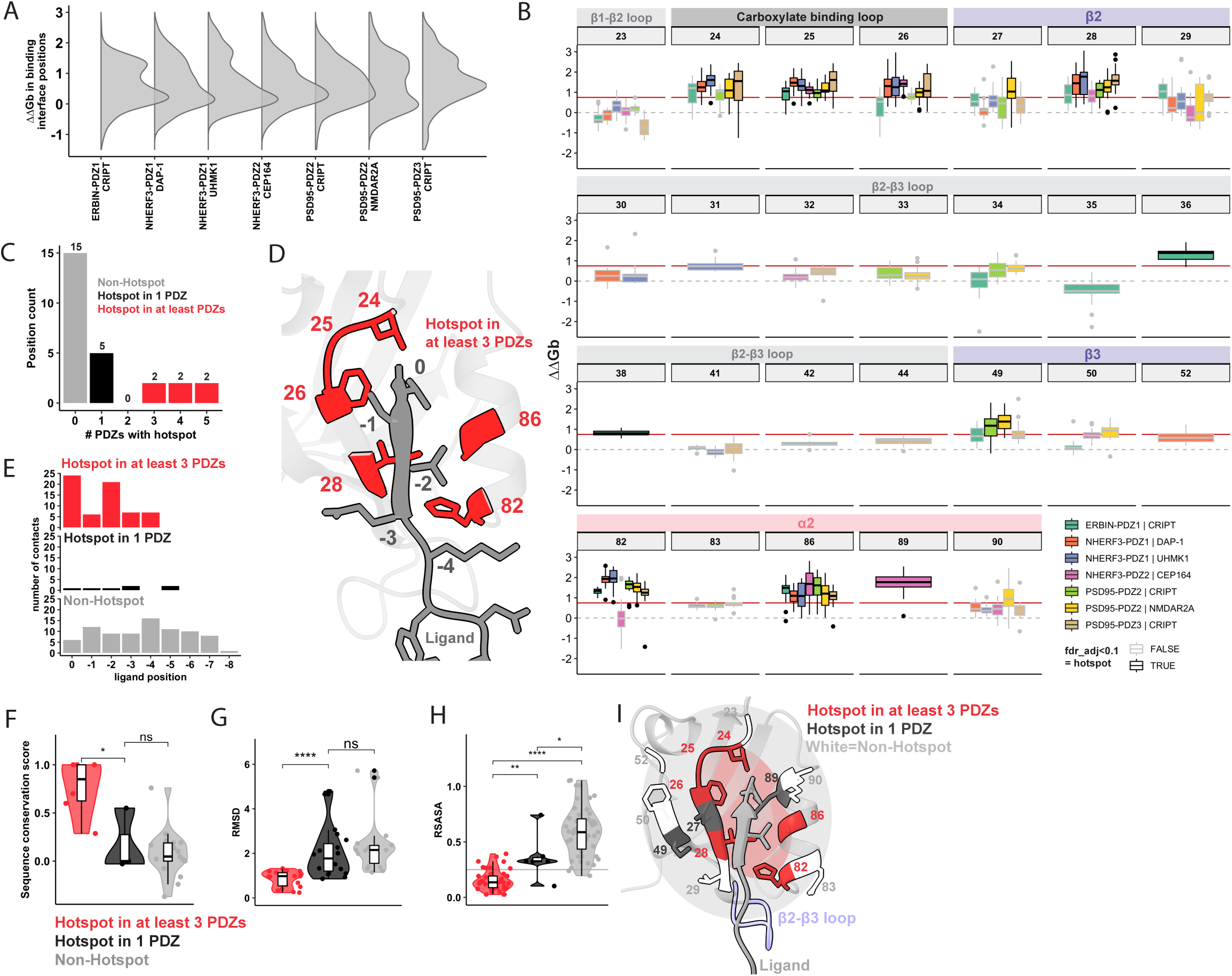
Comparative binding interface energy maps. **a.** Distribution of ΔΔG_b_ values for mutations in residues contacting each ligand. **b.** Distribution of ΔΔG_b_ values for mutations in each binding interface residue per interaction. Facets group structurally aligned residues from each interaction. The horizontal red line indicates the median ΔΔG_b_ across all the binding interfaces. Boxplots outlined in black highlight residues with significantly higher ΔΔG_b_ values than the median (ΔΔG_b_>0.75 kcal/mol, FDR < 0.1, one-sided t-test). **c.** Number of positions that are hotspots in 0 to 5 PDZ domains. Colors indicate non-hotspots (grey), hotspots in 1 domain (black), and recurrent hotspots in at least 3 (red). **d.** Structural view of hotspots conserved in at least 3 domains. **e**. Total number of PDZ residues contacting each ligand position. Facets separate contact counts from non-hotspots (grey), hotspots in 1 PDZ (black), and recurrent hotspots in at least 3 (red). **f**. Distribution of sequence conservation score per position in the alignment. “*” indicates *P*<0.05 (Wilcoxon rank-sum test). **g**. Distributions of Root-Mean-Square Deviation (RMSD) of residues within each group, computed for each pairwise comparison of binding interfaces. “*” indicate *P*<0.05 (Wilcoxon signed-rank test). **h**. Distribution of relative solvent accessible surface area (rSASA) per residue and interaction, grouped by hotspot classification. “*” indicate *P*<0.05 (Wilcoxon rank-sum test). **i**. Structural representation of the O-ring-like arrangement of hotspots in the binding interface. Residue labels correspond to structural alignment positions. Colors indicate non-hotspots (white), hotspots in 1 or 2 PDZ domains (dark grey), and recurrent hotspots in at least 3 PDZ domains (red).

Defining hotspots as residues where mutations cause changes in binding energy larger than the median of all interface residues (0.75 kcal/mol) identifies a median of six hotspots per interface (false discovery rate (FDR, Benjamini-Hochberg method) < 0.1, one-sided t-test). This is a median of 41.7% of interface residues (range 29.4% to 61.5% Fig. 3b). The interface hotspots are, however, not identical in all interactions: only two residues are hotspots in all seven interactions (structural alignment positions 25 in the carboxylate binding loop, and 86 in the α2 helix), and an additional four are conserved in more than half of the interactions (positions 24 in the carboxylate binding loop, 26 and 28 in β2, and 82 in α2, Fig. 3b,c,d).

Hotspots are enriched in contacts with ligand positions 0 or -2, the two aa positions used to define the three main classes of PDZ domain ligands (OR=13.71, *P*=2.61x10^-7^, one-sided Fisher’s Exact Test (FET) Fig. 3e)^17^. Indeed all six conserved hotspots contact at least one of these two ligand positions in all interactions, except Y24 in NHERF3-PDZ2 binding CEP164. The conserved hotspot positions also have higher sequence and structural conservation (Fig. 3f,g), and have highly conserved structural contacts, with five out of six positions contacting the ligand in all seven interactions (OR=60.54, *P*=5.3x10^-04^, one-sided FET).

The conserved hotspots are also buried deeper in the hydrophobic binding pocket (median rSASA = 0.14 vs. 0.33 for conserved and non-conserved hotspots, respectively, Wilcoxon rank-sum test *P=*5.8x10^-03^; median rSASA for non-hotspots=0.58, *P*=1.5x10^-14^, Fig. 3h). The hotspot residues occupy central positions at the interface, surrounded by a shell of energetically less important contacts across all protein interfaces analyzed (Fig. 3i, Extended Data Fig. 3a). This pattern is consistent with the ‘O-ring’ model of binding interface architectures, where surrounding residues exclude bulk solvent from central hotspots, enhancing binding affinity^28,30^. However, our data also show that residues in this outer ring can themselves evolve to become hotspots for certain interactions, for example positions 49 and 27 for PSD96-PDZ2, position 89 in NHERF3-PDZ2, and 36 and 38 in ERBIN-PDZ2 (Extended Data Fig. 3a,b).

### Interface hotspot evolution

A total of 15 interface residues are never classified as hotspots, including eight (positions 23, 29, 32, 34, 41, 50, 83, and 90) that make contacts with a ligand in multiple PDZ domains (Fig. 3b, extended Data Fig. 3a,c). These non-hotspot contacts are more solvent accessible (Fig. 3h) and they are particularly enriched in the highly variable β2–β3 loop^31^ (OR=6.23, *P*=0.04, one-sided FET, Fig. 3b and Extended Data Fig. 3a). However, residues in the β2-β3 loop can also function as hotspots. For example, positions G36 and P38 in the β2-β3 loop extension of ERBIN-PDZ1 add two additional protein-specific hotspots that recognise ligand position Y-5 (Extended Data Fig. 3a). These new hostposts outside of the canonical binding site contribute to a novel binding specificity by engaging non-canonical ligand positions^31^. The redistribution of binding energy to these new hotspots is accompanied by reduced energetic effects of mutations in the conserved residues L24, F26, and I28, resulting in a highly diverged interface energy landscape.

The energy landscape of NHERF3-PDZ2 binding a class 2 ligand is also highly diverged, with large energetic effects for mutations at V89, which contacts the class II ligand-defining phenylalanine ligand side chain at position 0 (Fig. 3b and Extended data Fig. 3a,b). The bulky F0 side chain requires small hydrophobic residues at V89 (such as V or I) (Extended data Fig. 3a,b). Once again, this new interaction is associated with reduced dependence on conserved hotspots L24, F26, and I28. Finally, two additional sites are only hotspots in PSD95-PDZ2: position S27 in the β2 strand and T49 in the β3 helix. These sites form non-canonical contacts with position -1 and -3, respectively.

### Seven comparative allosteric maps

We next focussed on changes in binding energy caused by mutations outside of the binding interfaces. These indirect effects on binding affinity are, by definition, allosteric^19,22,23^, allowing us to comprehensively compare allosteric communication in homologous proteins for the first time (Fig. 4a). In all five PDZ domains and for all seven interactions mutations outside of the binding interface can have strong effects on binding energy, (Fig. 4b), often comparable in magnitude to mutations in the binding interfaces (Fig. 4b,g).

**Figure 4:**
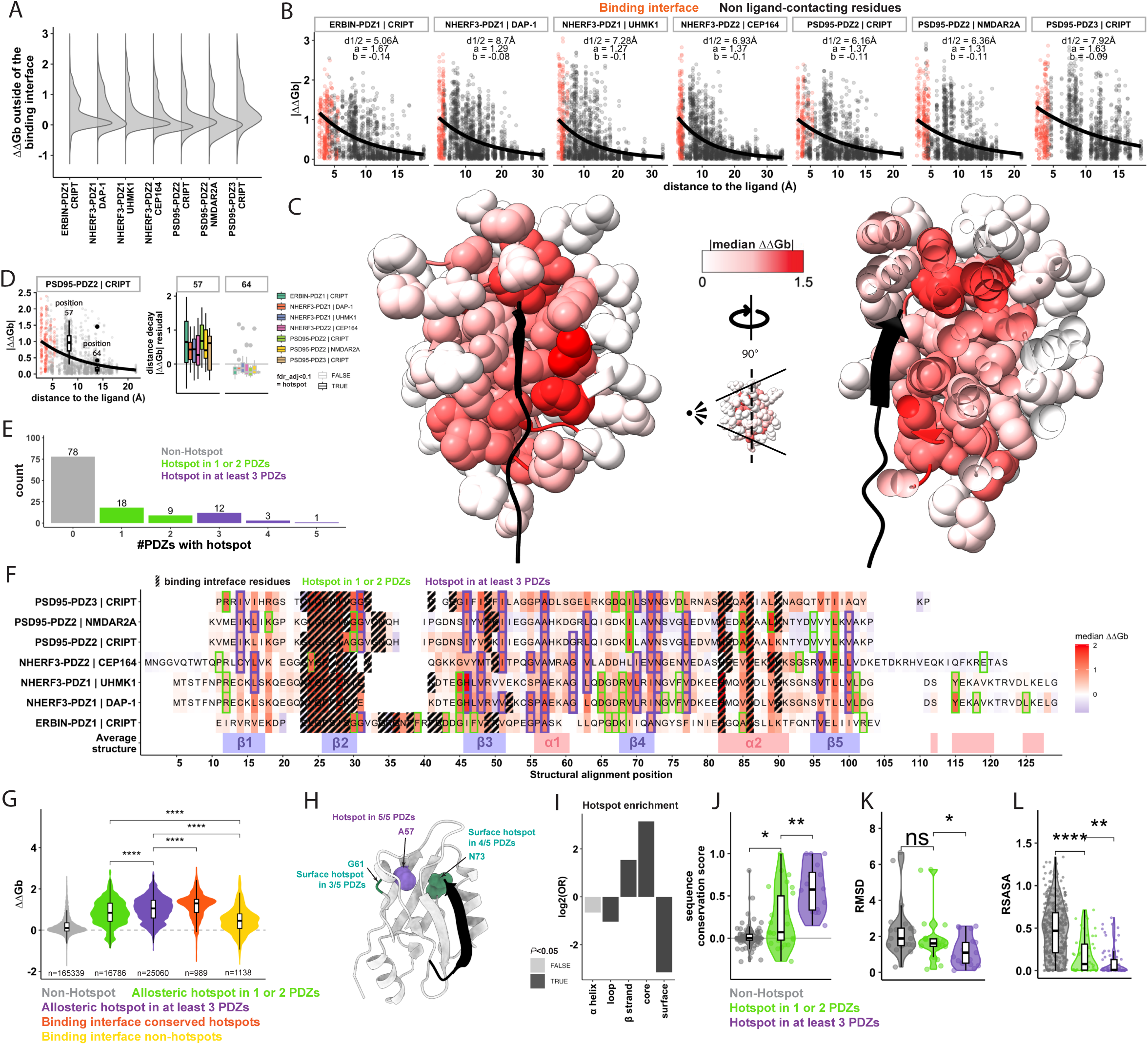
Seven comparative allosteric maps. **a.** Distribution of ΔΔG_b_ values for mutations in residues outside of the binding interface. **b.** Relationships between absolute ΔΔG_b_ and the minimum side chain heavy atom distance to the ligand. Curves represent exponential decay fits (*y = a · e^bx^*), calculated excluding mutations within the binding interface. **c.** Structural visualization of ΔΔG_b_ decay by distances to the ligand. ERBIN-PDZ1 binding CRIPT is shown, colored by absolute ΔΔG_b_ values. The second panel shows a sliced view highlighting internal residues. **d**. Examples of the ΔΔG_b_ (left) and ΔΔG_b_ residuals (right) distributions for structural alignment positions 57 and 63. Left panel shows the data for PSD95-PDZ2 binding CRIPT, and the right panel shows the distributions for all interactions. **e**. Number of positions that are allosteric hotspots in 0 to 5 PDZ domains. Colors indicate non-hotspots (grey), hotspots in 1-2 PDZs (green), and recurrent hotspots in at least 3 (purple). **f.** Structural alignment where residues are coloured by the per-residue median ΔΔG_b_. Binding interface residues are presented in black dashed lines. Green outlines indicate positions classified as hotspots in only one or two PDZ domains; purple outlines indicate conserved hotspots present in at least three PDZ domains. **g.** Distribution of ΔΔG_b_ values across five residue categories: non-hotspots outside the binding interface (gray), allosteric hotspots found in one or two PDZ domains (green), conserved allosteric hotspots found in three or more (purple), binding interface hotspots (orange), and binding interface non-hotspots (yellow). “*” indicate *P*<0.05 (Wilcoxon rank-sum test). **h.** Structure visualization of residues A57, G61, and N73. A57 is classified as an allosteric hotspot in all interactions tested. G61 is a surface-exposed allosteric hotspot in 3 out of 5 PDZ domains, and N73 is a surface-exposed allosteric hotspot in 4 out of 5 PDZ domains. **i**. Two-sided Fisher’s exact test for enrichments of allosteric hotspots in alpha helices, loops, beta strands, and One-sided Fisher’s exact test for enrichment in core (rSASA<0.25) and surfaces. Tests are based on the complete dataset aggregated from all libraries. **j**. Distribution of sequence conservation score per position in the alignment. “*” indicates *P*<0.05 (Wilcoxon rank-sum test). **k**. Distributions of Root-Mean-Square Deviation (RMSD) of residues within each group, computed for each pairwise comparison of structures. “*” indicates *P*<0.05 (Wilcoxon signed-rank test). **l**. Distribution of relative solvent accessible surface area (rSASA) per residue and interaction, grouped by hotspot classification. “*” indicate *P*<0.05 (Wilcoxon rank-sum test).

### Conserved distant-dependent allosteric decay

We first considered how the effects of mutations on binding energy relate to their distance to the ligand. Plotting absolute ΔΔG_b_ against. the Euclidean distance to the nearest ligand residue reveals a strikingly conserved relationship across all seven interactions: the probability of a mutation causing a change in binding energy decays with the distance from the binding interface, with a 50% reduction of allosteric effects over a median distance d1/2 = 6.9 Å (Fig. 4b,c).

This decay is similar to that quantified for allostery in the SRC kinase domain^23^ (d1/2 = 8.3 Å, Extended Data Fig. 4a), KRAS for six interaction partners^22^ (median d1/2 = 9.50Å, Extended Data Fig. 4b), GB1^19^ (d1/2 = 5.6 Å), and GRB2-SH3 domain^19^ (d1/2 = 13.6 Å, Extended Data Fig. 4c). This conserved distance-dependent decay of allosteric efficacy in nine different proteins suggests it is a general principle of allosteric communication (Fig. 4c).

### Allostery beyond distance-dependent decay

We next quantified the allosteric effect of each mutation beyond that expected from its distance to the ligand. For each mutation we calculated the residual between its measured change in binding energy, ΔΔG_b_, and that expected given the fitted distance-dependent decay (Fig. 4b). These ΔΔG_b_-residuals quantify how much more allosteric each mutation is than expected given the Euclidean distance to the ligand.

The ΔΔG_b_-residuals for mutations outside of the binding interfaces are well correlated for the same PDZ domains binding to two different ligands (median Pearson’s *r* = 0.81) but are less conserved when comparing between two different PDZ domains (median *r* = 0.50, Extended Data Fig. 4d). For each interaction, a median of 14.4% of mutations have effects larger than expected given the distance-dependent decay (one-sided Z-test for residual>0, *FDR*<0.1). Across pairwise comparisons, only a median of 7.3% of these unusually strong allosteric mutations identified in one interaction are also classified as unexpectedly allosteric in a second PDZ domain. However, the overlap for the same PDZ domain binding different ligands is larger, increasing to a median of 41.4% (Extended Data Fig. 4e).

### Allosteric hotspots

We define allosteric hotspots as positions where mutations cause larger changes in ΔΔG_b_ than expected at their distance from the ligand (*FDR*<0.05, one-sided t-test; median |ΔΔG_b_| > 0.2kcal/mol) (see methods, Fig. 4d,f and Extended Data fig. 4f). For each interaction, this defines a median of 19.6% of the positions outside the binding interface as allosteric hotspots (range 17.4% to 24.7%). Mutations in these allosteric hotspots cause changes in binding energy comparable in magnitude to mutations in the binding interface (Fig. 4g and Extended Data Fig. 4g).

Across the five proteins, 43 different positions are allosteric hotspots for at least one interaction while 78 positions are not hotspots for any interaction (Fig. 4e). One position (A57 in α1 helix) is an allosteric hotspot in all seven PDZ domain interactions tested (Fig. 4e,h & Extended Data fig. 4f). Position A57 contacts the carboxylate-binding loop in all interactions and was previously reported as allosteric in PTP-BL-PDZ2 (A46) and PSD95-PDZ3 (A347)^32^. 16 residues are hotspots in three or more domains, nine are hotspots in two domains, and 18 are only hotspots in a single protein (Fig. 4e). The hotspots are enriched in β-strands (OR=2.90, *P*=1.01x10^-06^, two-sided FET) and depleted in loops (OR=0.49, *P*=5.88x10^-04^, Fig 4i). The allosteric hotspots for PSD95-PDZ3 strongly overlap with residues reported and predicted as allosteric in previous experimental and computational studies (Extended Data Fig. 4h).

### A conserved allosteric core

The 43 allosteric hotspot residues are strongly enriched in the previously described ‘sector’ of co-evolving residues in PDZ domains^33,34^ (n=11, OR=26.13, *P*=2.96x10^-05^, one-sided FET considering all residues outside of the binding interface), and this enrichment is stronger for the 15 most conserved hotspots (OR=30.02, *P*=3.64x10^-06^). Similarly, the most conserved six binding interface hotspots are also enriched in the co-evolving sector (OR=17.16, *P*=9.6x10^-03^). The complete set of allosteric hotspots is also enriched in the protein core (rSASA<0.25, OR=8.91, *P*<2.2x10^-16^, one-sided FET, Fig. 4i, extended Data Fig.4i). Indeed 14 of the 16 most conserved allosteric hotspots are always classified as buried in the core in domains where they are classified as hotspots (OR=13.77, *P*<2.2x10^-16^), with the only exceptions being surface residues at positions 73 and 61 (Fig. 4h). The most conserved hotspots are also more likely to be core residues compared to hotspots only classified in one or two PDZ domains (OR=3.35, *P*=1.10x10^-02^, Fig. 5a-g). The most functionally conserved allosteric hotspots are also more conserved in sequence and structure (Fig. 4j,k).

**Figure 5:**
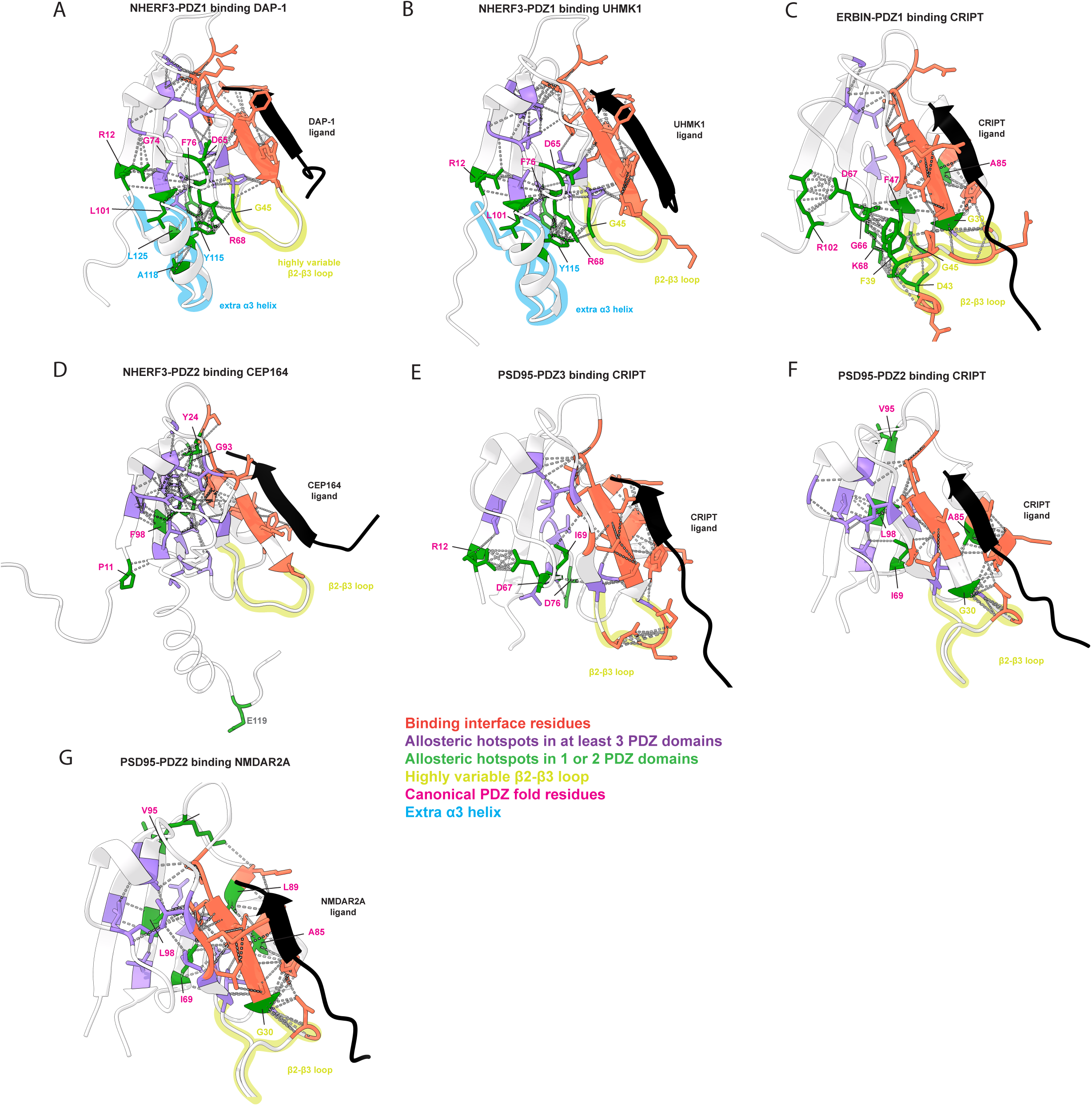
Allosteric innovation in a protein family. Binding interface residues (ligand-contacting) are shown in red. Allosteric hotspots present in at least 3 PDZ domains are shown in purple; those found in only 1-2 PDZs are shown in green. Non-conserved hotspots located in canonical PDZ fold regions are labelled in pink. β2-β3 loop regions are shown in yellow, and additional C-terminal α-helix extensions are in blue. Predicted direct contacts between hotspots are indicated with grey dashed lines. Structure for **a.** NHERF3-PDZ1 binding DAP-1 **b**. for NHERF3-PDZ1 binding UHMK1 **c**. ERBIN-PDZ1 binding CRIPT **d**. NHERF3-PDZ2 binding CEP164 **e**. PSD95-PDZ3 binding CRIPT **f.** PSD95-PDZ2 binding CRIPT **g.** PSD95-PDZ2 binding NMDAR2A.

The PDZ domains therefore contain a conserved allosteric core of co-evolving residues. However, each individual protein also contains additional allosteric residues. We turn to these in the next section.

### Allosteric innovation in a protein family

In addition to the functionally conserved hotspots, for each interaction there is a median of seven additional hotspots only present in one or two PDZ domains. Binding energy changes in these more protein-specific hotspots are well-correlated for the same PDZ domain binding to different ligands (Pearson’s *r* = 0.93 for PSD95-PDZ2 and *r* = 0.71 for NHERF3-PDZ1, Extended Data Fig. 4j,k) but are less correlated when comparing different proteins binding the same ligand (median *r* = 0.45) or different proteins binding to different ligands (median *r* = 0.32, Extended Data Fig. 4l). We next examined the structural location of these less conserved hotspots.

In NHERF3-PDZ1 the less conserved hotspots are spatially clustered in a region that contains an insertion of an extra α-helix (Fig. 5a,b). Domain insertions containing additional secondary structure elements are quite frequent in PDZ^11^ and other^10^ domains and NHERF3-PDZ1 illustrates how these insertions can connect to the conserved allosteric network. Interestingly, this same region contains a spatial cluster of novel hotspots in ERBIN-PDZ1, despite this domain not containing an additional α-helix (Fig. 5c). The domain does, however, contain multiple aa insertions in the β2-β3 loop in this region. The β2-β3 loop of ERBIN-PDZ1 contains two protein-specific hotspots and one hotspot shared with NHERF3-PDZ1.

The spatial cluster of novel allosteric hotspots in NHERF3-PDZ1 and ERBIN-PDZ1 link the binding interface to the opposite surface of the domain (Fig. 5a,b and 6b). ERBIN-PDZ1 has the most distinct set of allosteric hotspots of any of the domains, with 10 novel hotspots shared with at most one other PDZ domain and only five hotspots shared with three or more domains (compared to a median of 10 for the other interactions). Thus, the loop extension in this domain appears to have expanded and partly replaced the typical allosteric network.

Not all domain extensions are, however, strongly coupled to the core allosteric network. NHERF3-PDZ2 contains a 21 aa C-terminal extension but mutations in only one residue (E119) have larger effects on binding than expected (Fig. 5d), with a median ΔΔG_b_ = -0.20 kcal/mol. (Note that this extension is likely disordered, see methods, and may potentially dynamically directly contact the ligand rather than being allosteric, so we conservatively do not classify it as an allosteric hotspot.)

Other protein-specific hotspots in the PDZ domains are structurally conserved sites, so the mechanisms underlying their allosteric evolution are less clear. For example, in PSD95-PDZ3, a less conserved set of hotspots extends the conserved allosteric core to the protein surface opposite to the binding interface (surface residues R12, N73 and D76, Fig. 5e, Fig. 6b). In contrast, for both interactions of PSD95-PDZ2, most of the novel allosteric hotspots are buried in the protein core (G30, I69, A85, L89 and L98) with two novel hotspots on the surface (K18 and V95).

**Figure 6:**
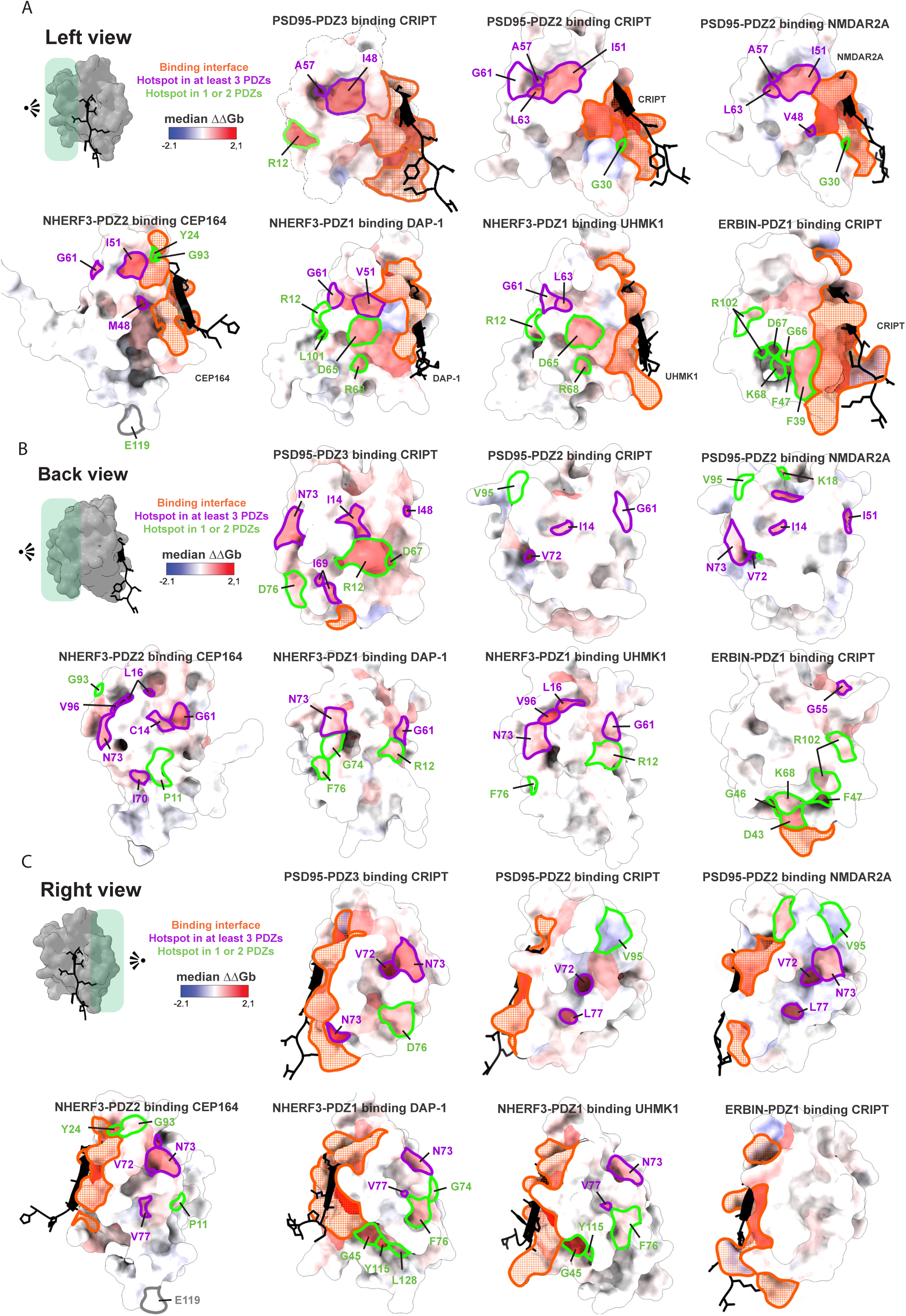
Evolution of allosteric surfaces. Surfaces are colored by the per-residue median ΔΔG_b_. The binding interface is highlighted in orange. Residues outlined in purple are allosteric hotspots present in at least three PDZ domains; green outlines indicate hotspots identified in only one or two domains. The allosteric surface map is shown from three orientations relative to the reference structure, viewed facing the ligand with the C-terminus pointing upward: **a**. from the left, **b**. from the back, **c**. from the right.

### Gain-of-function allosteric mutations

Across all five domains and seven interactions there is only one allosteric hotspot where mutations on average increase ligand binding (median ΔΔG_b_<0): position V95 of PSD95-PDZ2 (Extended Data Fig. 4f). V95 is solvent-exposed and located in the β5 strand and all mutations increase binding (ΔΔG_b_<0, *FDR* < 0.1, Z-test), with a median ΔΔG_b_ of -0.45 kcal/mol for binding to CRIPT and -0.45 for binding to NMDAR2A (Fig. 5f,g).

Interestingly, the V95 surface site in PSD95-PDZ2 does not directly contact any other hotspot residues for binding to CRIPT and contacts only one hotspot for binding to NMDAR2A (L16).

To further investigate the characteristics of GOF mutations we considered all mutations in all domains causing a decrease in binding energy (ΔΔG_b_<0). The distance-dependent decay for these GOF mutations is weaker than for mutations that are detrimental for binding (median decay rate b=-0.054 and d_1/2_=11.89 Å for ΔΔG_b_<0, and b=-0.096, d_1/2_=7.23 Å for ΔΔG_b_>0) (Extended Data Fig. 5a). Interestingly, GOF allosteric mutations in the Src protein kinase domain also show a much weaker distance-dependence than loss-of-function mutations^23^.

To further investigate the spatial location of GOF mutations we considered all mutations in all domains causing a strong decrease in binding energy (ΔΔG_b_ < -0.2 kcal/mol and FDR < 0.1, one-sided Z-test for ΔΔG_b_<0). Across domains, between one (PSD95-PDZ3) and 19 (PSD95-PDZ2 binding NMDAR2A) residues are enriched for these GOF mutations (FDR<0.1, one-sided FET, Extended Data Fig. 5b), including multiple surface and in loop residues (Extended Data Fig. 5b,c), but their location is dispersed and varies across PDZ domains and interactions (Extended Data Fig. 5c). GOF mutations in PSD95-PDZ2 do, however, show spatial clustering in the β4 strand, β4-α2 loop, and α2-β5 loop, regions that are close to the V95 GOF allosteric hotspot (Extended Data Fig. 5).

The dynamic range of our selections provides less power to detect GOF variants than variants that decrease binding. However, their spatial architecture does seem different, as has previously been reported for Src^23^, with the strong protein-specific GOF allosteric hotspot in PSD95-PDZ2 illustrating how surface-exposed sites can evolve to have strong positive effects on binding when perturbed.

#### Evolution of allosteric surfaces

Finally we considered the accessible allosteric surfaces of the five homologous domains. Surface sites are particularly important for regulation and also provide accessible sites to be targeted by therapeutics^4,22,23,35^. We therefore more comprehensively compared the allosteric surfaces of the five PDZ domains (Fig. 6). In all five PDZ domains there are at least two solvent-exposed allosteric hotspots, with between 4.3% and 9.1% of the surface residues defined as hotspots (median of 7.3%) and a total of 14 unique solvent accessible hotspots. Two surface positions are hotspots in a majority of domains: positions 61 and 73 in the structural alignment (Fig. 4h). Position 61 is a hotspot for 4/7 interactions and is a glycine in all domains except ERBIN-PDZ1 where it is deleted. G61 is located in the α1-β4 loop, a median of 13.94 Å from the ligand and mutations in this residue cause a median ΔΔG_b_=0.70 kcal/mol (Fig. 6a,b). Position 73 is a hotspot for 5/7 interactions and is located in the β4-α2 loop, at a median distance of 11.56 Å from the ligand (median ΔΔG_b_ = 0.86 kcal/mol, Fig. 6b,c). Changes in binding energy (ΔΔG_b_) for mutations in surface hotspot positions correlate strongly for the same domain binding to different ligands (median Pearson’s *r* = 0.87), but more weakly between different PDZ domains (median *r* = 0.23), further emphasising the protein-specificity of surface allostery.

Approximating PDZ domains as a cuboid, all three of the largest faces - the ‘left’, ‘right’ and ‘back’ surfaces with respect to the binding interface - contain allosteric hotspots in multiple domains (Fig. 6). However, the allosteric surfaces differ quite extensively across the five domains, indicating again that the peripheral allostery is quite fast evolving.

Considering the ‘left’ surface, the allosteric map is quite similar for PSD95-PDZ2 and PSD95-PDZ3, with a small allosteric pocket containing I51, A57 and L63 (Fig. 6a). However, PSD95-PDZ3 also contains an additional protein-specific hotspot, position R12. I51 is also an allosteric hotspot in NHERF3-PDZ2, but its context has changed (Fig. 6a). For NHERF3-PDZ1 the left allosteric surface has extended ‘down’ in the direction towards more N-terminal ligand residues, and in ERBIN-PDZ1 allosteric residues are now concentrated in this lower region including a pocket created by the β2-β3 loop extension (Fig. 6a).

The ‘back’ surface opposite the binding interface is less allosterically active in most of the domains. However, PSD95-PDZ3 contains a novel strong allosteric hotspot on this surface, residue R12, that forms a salt bridge with D67 (Fig. 6b). Other allosteric hotspots visible on the back surface have much smaller solvent accessibility and/or small median mutational effects (Fig. 6b).

Considering the ‘right’ surface (Fig. 6c), position N73 is a highly conserved allosteric hotspot. This residue is located on the surface in all interactions. In addition, in three PDZ domains - PSD95-PDZ2, PSD95-PDZ3, and NHERF3-PDZ2 - the adjacent and less solvent-accessible residue, V72, is also allosteric. NHERF3-PDZ1 has evolved two additional allosteric patches lower down the right surface (residues F76, G45, Y115, and L128). Residue 76 in the lower right surface is also allosteric in PSD95-PDZ3. In contrast, PSD95-PDZ2 and NHERF3-PDZ2 have additional hotspots towards the top of the right surface, with V95 in PSD95-PDZ2 the only positive allosteric hotspot across all domains. Finally, ERBIN-PDZ1 contains no allosteric hotspots on the right surface with mutations across the entire surface having very limited effects on binding energy. Indeed in ERBIN-PDZ1 the β2-β3 loop extension appears to have shifted the allosteric network ‘down’ and ‘left’.

In summary, all five PDZ domains contain multiple allosteric surface sites but this surface allostery appears to be quite fast evolving. The shift of allostery ‘up’ and ‘down’ or ‘left’ and ‘right’ in the homologous domains means that each has a unique allosteric surface for cellular regulation and a unique allosteric surface to therapeutically target.

## Discussion

We have presented here the first comprehensive energetic and allosteric maps for homologous proteins. Using pooled selection and sequencing experiments we quantified 424,512 changes in abundance and 619,072 changes in binding for single and double mutant variants of five human PDZ domains interacting with seven different ligands, allowing us to infer the changes in free energy of folding and binding for all amino acid substitutions, a total of 21,802 free energy measurements. The resulting data provide a number of important insights into the evolution of allostery.

First, despite their small size, all five PDZ domains are allosteric and all five show a strong distance-dependent decay of allostery away from the binding interface. This distance-dependent decay is also observed in all four published experimental allosteric maps^19,22,23^ (Extended Data Fig. 4a,b,c), for energetic couplings in molecular dynamics simulations^36^, for evolutionary conservation around enzyme active sites^37^, and for chemical shift perturbations in NMR experiments^36^. We believe therefore it is likely to be a general and conserved principle of protein biophysics.

Second, our data reveal a modular allosteric architecture of PDZ domains, with a conserved allosteric core connecting to protein-specific extensions to surface sites. This modular architecture is also highly consistent with evolutionary analyses, which have revealed a conserved co-evolving core (or ‘sector’) of PDZ and other domains^33,34,38,39^, and experiments identifying allosteric surface sites connected to co-evolving sectors^39,40^. Our data suggest that allostery primarily evolves via the gain-and-loss of peripheral extensions to a conserved allosteric core. It will be interesting in future work to test if this conservation of an allosteric core with faster evolution of the allosteric surface is also observed in other protein families.

Third, we believe the faster evolution of the allosteric surface has important implications for drug development. In each protein a median of 7.3% of solvent-exposed surface sites are allosterically active and so potentially targetable. However, the most allosteric surface sites change across the five proteins, suggesting that the best sites to engage therapeutically differ in different members of the protein family. If this principle of fast evolving allosteric surfaces extends to other protein families it should allow the development of both more allosteric drugs and more specific allosteric drugs that target one protein without inhibiting or activating other family members, even if they are bound. The faster structural evolution of surface sites compared to orthosteric sites^5,6^ provides another mechanism to increase specificity. It will be important in future work to test this hypothesis in other large protein families, including those of high therapeutic value such as protein kinases and G-protein coupled receptors (GPCRs).

Our dataset also provides – to our knowledge – the first complete comparative energy maps for homologous protein interaction interfaces. Most of the hotspot residues important for binding are conserved across the PDZ domains. However, additional hotspots have evolved, including by the insertion of amino acids into a loop. In future work it will be important to extend this comparative mapping of binding interface energy landscapes to additional protein families.

We believe comprehensive comparative energy landscapes such as those presented here are important for understanding both protein evolution and how to therapeutically target proteins from large families. Moreover, scaling the production of energy maps should produce well-calibrated datasets that will allow the full potential of machine learning to be brought to bear on biological discovery, drug development, and biotechnology.

Targeting allosteric sites has enormous potential for the development of therapeutics, allowing the development of inhibitors, activators and modulators of proteins previously considered ‘undruggable’ and also allowing the development of more specific and less toxic treatments^1,4,35^. Moreover, engineering allosteric control is important in biotechnology, facilitating the engineering of biosensors, control circuits and metabolic pathways^41–43^. The modular allosteric architecture that we have identified here for PDZ domains further reinforces this potential for targeting and engineering allostery across a protein family. In future work it will be important to explore the comparative allosteric architectures of additional protein families and functions, including those of high therapeutic and technological value such as receptors, enzymes, and transcription factors.

## Methods

### Media

- LB: 10 g/L Bacto-tryptone, 5 g/L Yeast extract, 10 g/L NaCl. Filter sterilised
- YPD: 20 g/L glucose, 20 g/L Peptone, 10 g/L Yeast extract.
- YPDA: 20 g/L glucose, 20 g/L Peptone, 10 g/L Yeast extract, 40 mg/L adenine sulphate. Filter sterilised.
- SORB: 100 mM LiOAc, 10 mM Tris pH 8.0, 1 mM EDTA, 1 M sorbitol
- Plate mixture: 100 mM LiOAc, 10 mM Tris-HCl pH 8.0, 1 mM EDTA pH 8, 40% PEG3350. Filter sterilised.
- Recovery medium: YPD + Sorbitol 0.5M. Filter sterilised.
- SC -URA: 6.7 g/L Yeast Nitrogen base without amino acid, 20 g/L glucose, 0.77 g/L complete supplement mixture drop-out without uracil. Filter sterilised.
- SC -URA/ADE: 6.7 g/L Yeast Nitrogen base without amino acid, 20 g/L glucose, 0.76 g/L complete supplement mixture drop-out without uracil, and adenine. Filter sterilised.
- SC -URA/MET/ADE: 6.7 g/L Yeast Nitrogen base without amino acid, 20 g/L glucose, 0.74 g/L complete supplement mixture drop-out without uracil, adenine and methionine. Filter sterilised.
- MTX competition medium: SC -URA/ADE + 200 ug/mL methotrexate (BioShop Canada Inc., Canada), 2% DMSO.
- DNA extraction buffer: 2% Triton-X, 1% SDS, 100mM NaCl, 10mM Tris-HCl pH8, 1mM EDTA pH8.

### Wild-type PDZ domain testing

To design the libraries used, we first tested 20 PDZ wild-type sequences, 4 from Salinas and Ranganathan 2018^26^, and 16 from Teyra et al. 2020^45^. PDZ domain boundaries were manually defined, taking into account those used in Teyra et al. 2020^45^, and available PDB structures, ensuring that secondary structure elements were not truncated or excluded. These sequences were tested for signal in AbundancePCA^19^. Of the 13 wild type PDZ domains that showed signal, we further tested 24 interactions in BindingPCA. These included: natural ligands, top-scoring natural proteins identified from optimal motif matches in Teyra et al. 2020^45^, and the CRIPT ligand tested across several of the assayed PDZ domains. Ligand sequences include the C-terminal 9 amino acids of the interaction proteins.

The domain and ligand wild type nucleotide sequences were obtained from a custom randomised back translation algorithm to increase codon complexity, which also avoided the restriction sites used in the cloning process: NheI, HindIII, SpeI and BamHI (see below). Gibson complementarity regions were fused to the extremes of the designed sequences. Common 5’ acatttccccgaaaagtggggaggtggagctagc and 3’ taaaagcttcgcaggaaagaacatgtgagcaaaa for PDZ domains. Common 5’ aaaagtgactagttta and 3’ ggatccgcaatgtaaa for ligand sequences. These added regions also introduced NheI, and HindIII for PDZ sequences, and SpeI and BamHI for ligand sequences (see below). Designed PDZ wild type sequences oligos were ordered through TWIST and ligand sequences through IDT as an oligo pool.

Three main plasmids were used to clone the wild type PDZ sequences; the two generic plasmids for AbundancePCA (pGJJ001) and BindingPCA (pGJJ045) as in^19^, and a generic mutagenesis plasmid (pGJJ191), which contained a streptomycin resistance gene cassette.

Ligands designed for each interaction were inserted into the target BindingPCA plasmid by using a two-step cloning strategy. All single stranded ligand sequences, obtained in a pool, were amplified using oGJJ789 primer in one single PCR cycle. pGJJ191 empty backbone was linearised using oGJJ787 and oGJJ788 primers, which introduced SpeI and BamHI restriction sites (all the primers used are described in Supplementary Table 1). The pool of ligand sequences was then inserted into the pGJJ191 vector by a Gibson (NEB) reaction at 50°C for 3h. Single transformants were individually checked by Sanger sequencing and correct ones were later cloned in separate reactions into the binding vector pGJJ001 by digestion ligation using the SpeI and BamHI restriction enzymes and T4 ligase (NEB). Digested ligand sequences were gel purified using minElute gel extraction kit (QIAGEN). Empty digested vector pGJJ001 was verified through gel electrophoresis and purified using QIAquick PCR purification kit (QIAGEN). This reaction cloned the ligand N-terminally fused to DHFR3.

The wild-type sequences of the several PDZs were amplified using a common reverse PCR primer oGJJ718, and assembled by Gibson (NEB) reaction at 50°C for 3h into the pGJJ191 vector previously linearised using oGJJ728 and oGJJ734 primers, which introduced NheI and HindIII restriction sites. Later, PDZ wild type sequences from pGJJ191, the empty pGJJ045 abundance and the already-containing ligand pGJJ001 binding vector were digested using NheI and HindIII restriction enzymes. Digested PDZ sequences were gel purified using minElute gel extraction kit (QIAGEN). Empty digested vectors (pGJJ045 and pGJJ001) were verified through gel electrophoresis and purified using QIAquick PCR purification kit (QIAGEN). Final digested wild-type PDZ sequences were assembled into the empty pGJJ045 abundance and the already-containing ligand pGJJ001 binding vector using T4 ligase (NEB). This reaction cloned the ligand N-terminally fused to DHFR1,2.

At each step, gibson products were dialyzed, concentrated to 5μL using a SpeedVac machine, and transformed into NEB 10β competent *E. coli* cells according to the manufacturer’s protocol. Cells were recovered in SOC (NEB 10β Stable Outgrowth Medium) for 30 minutes and plated for later DNA extractions from single colonies. Correct plasmid construction was verified at each step using Sanger sequencing (GATC, Eurofins Genomics) from mini prep products.

Cloned plasmids were then individually transformed into yeast using a small scale yeast transformation standard protocol as in Faure et al. 2022^19^. After 48 hours, transformed cells were subject to selection in the presence of methotrexate (see below). By measuring the optical densities at 600nm (OD600) every 15 minutes for a minimum of 60 hours, we calculate the growth curve under selection for each transformation. Half-growth times were calculated as the time required for the culture to reach half of its maximum OD600. Domains with mean half-growth time lower than 40 hours in both AbundancePCA and their interactions in BindingPCA were considered for further experimental mutagenesis.

### Library designs

From the tested wild types, we finally designed mutagenesis libraries on 4 different PDZ domains (PSD95-PDZ2, NHERF3-PDZ2, NHERF3-PDZ1 and ERBIN-PDZ1) that were proven to grow in abundance and binding assays against several interaction partners. This set of libraries included also the following PDZ interactions: PSD95-PDZ2 binding CRIPT (Uniprot IDs: P78352 and Q9P021, respectively), PSD95-PDZ2 binding NMDAR2A (P78352 and Q12879, alternative name NR2A), NHERF3-PDZ2 binding CEP164 (Q5T2W1 and Q9UPV0), NHERF3-PDZ1 binding DAP-1 (Q5T2W1 and O14490), NHERF3-PDZ1 binding UHMK1 (Q5T2W1 and Q8TAS1), and ERBIN-PDZ1 binding CRIPT (Q96RT1 and Q9P021) (Supplementary Table 2). We did this in addition to the already existing data for PSD95-PDZ3 binding CRIPT (P78352 and Q9P021) from Faure et al. 2022^19^. This makes a final dataset of 5 PDZs assayed in AbundancePCA and a total of 7 interactions tested in BindingPCA.

Those libraries were splitted into two blocks, for the 1st and 2nd halves of the PDZ sequences respectively (Fig. 1C). The two blocks mutagenise overlapping parts of the sequences for further normalization of results (see below). The overlapping region between blocks included 5 amino acids in each case. Each library consisted of an oligo pool ordered through IDT where each codon was mutagenised to the degenerate NNK codon, to include all possible mutations in each position, and together with 32 genetic backgrounds, also codified as an additional fixed NNK position. Final oligos were a maximum length of 220 nucleotides.

### Plasmid library construction

This first step consisted of introducing the mutagenised oligos for each library block (ordered as oligo pools through Integrated DNA Technologies, IDT) into the mutagenesis plasmid already containing the wild-type version of the protein. This step is needed to build the mutagenesis library on each half of the protein sequence, whilst maintaining the other half untouched. Block2 libraries were amplified using the common oGJJ718 primer. Each of the block1 libraries was amplified using a distinct PCR primer, specific for each PDZ constant region, as shown in Supplementary Table 1. Library amplification was performed in 10 cycles for block1 libraries, and 1 PCR cycle for block2 libraries. Amplified products were purified using MinElute PCR purification kit. Plasmid mutagenesis backbones (PDZ wild-type containing pGJJ191) were also linearised using different sets of primers for each library’s constant regions, as shown in Supplementary Table 1. Primers annealing to constant regions outside of the PDZ sequences were common for each block of libraries; common reverse primer oGJJ891 for the block1 libraries, and common forward primer pGJJ720 for the block2 libraries. In separate reactions, each library was inserted by Gibson (NEB) reaction at 50°C for 3h into the PCR linearised pGJJ191 containing the other half of the wild type PDZ sequence. Gibson reactions were performed using in-house made master mix and a 1:5 vector-to-library ratio was used. Gibson products were dialyzed, concentrated to 10μL using a SpeedVac machine, and transformed into NEB 10β High-efficiency Electrocompetent *E. coli* cells according to the manufacturer’s protocol. Cells were recovered in SOC medium (NEB 10β Stable Outgrowth Medium) for 30 minutes. From the recovered product, 1μL of transformed cells were plated into LB spectinomycin plates to estimate the number of transformants. A minimum 20x coverage was aimed at each transformation, as shown in Supplementary Table 3. The remaining volume was transferred to 200mL of LB with spectinomycin to grow overnight. 50mL of grown culture was collected the next morning for DNA extraction using the QIAfilter Plasmid Midi Kit (QIAGEN).

Second step of this cloning strategy consisted of introducing the whole protein sequences (with half of their sequence mutagenised) into the AbundancePCA (pGJJ045) and BindingPCA plasmids (pGJJ001 already containing the corresponding ligand). Extracted library-containing plasmids were digested at NheI and HindIII restriction sites. Digested libraries were gel purified using Minelute gel extraction kit (QIAGEN), and digested plasmids were verified through gel electrophoresis and purified using QIAquick PCR purification kit (QIAGEN). Purified products were assembled using an overnight temperature-cycling ligation reaction with T4 ligase (NEB). As in the previous step, products were transformed into NEB 10β High-efficiency Electrocompetent *E. coli* cells according to the manufacturer’s protocol. A minimum 20x coverage was aimed at each transformation (Supplementary Table 3). Cells were recovered in SOC medium (NEB 10β Stable Outgrowth Medium) for 30 minutes. From the recovered product, 1μL of transformed cells were plated to estimate the number of transformants, and the remaining volume was transferred to 200mL of LB with ampicillin to grow overnight. 50mL of grown culture was collected the next morning for DNA extraction using the QIAfilter Plasmid Midi Kit (QIAGEN). Cloned libraries at each step were verified using oxford nanopore sequencing (Plasmidsaurus and Source Biosciences).

### Library pooling

Library selections, including AbundancePCA and BindingPCA libraries, were pooled based on the time needed to reach half OD600. Libraries with similar growth times were combined into the same competition assay to ensure proper selection at the output harvesting time. Since each PDZ was present in more than one library (e.g. in at least two assays), library groups were arranged to ensure that no PDZ sequence was repeated within the same pooling group, as shown in Supplementary Table 3. This arrangement ensured proper separation of libraries in later sequencing steps.

### Large-scale transformations of libraries into yeast

Each pool of libraries was transformed in triplicate using a high-efficiency yeast transformation protocol as described in^19^. The protocol described below (adjusted to a pre-culture of 1L of YPDA) was scaled up or down in volume depending on the targeted number of transformants of each library pool, as reported in Supplementary Table 3.

For each selection assay, three independent pre-cultures of BY4742 (Saccharomyces cerevisiae, MATα his3Δ1 leu2Δ0 lys2Δ0 ura3Δ0) were grown in 100 ml standard YPDA at 30°C overnight. The next morning, three replicates of 1L of YPDA were inoculated at optical density at OD600 of 0.3. After incubation at 30°C for 4h, the cells were collected and centrifuged for 5 min at 3,000g, washed with sterile water, and later with SORB medium, the cells were resuspended in 43ml of SORB and incubated 30 minutes at room temperature on a rotating wheel. Later, 875μL of 10 mg/mL boiled salmon sperm DNA (Agilent Genomics) was added to each tube of cells, along with 17.5 μg of equimolarly pooled plasmid libraries, pooling volumes adjusted based on both library complexity and DNA concentration. Then, 175 mL of plate mixture were added and incubated at room temperature for 30 minutes on a rotating wheel. After incubation, 17.5mL of DMSO were added and each sample was splitted into 5 equal 50mL falcon tubes. After gently mixing, samples were heat shocked at 42 °C for 20 min while gently inverting the tubes regularly. Later, cells were pooled again into the previous 1L rotor tubes to centrifuge for 5 minutes at 3000rpm. Supernatant was carefully removed, cells were resuspended in 250mL of recovery media and incubated at 30 °C for 1h. Once recovered, cells were again centrifuged 5 minutes at 4000rpm, and resuspended in 1L of plasmid selection media SC-URA. From this sample, 10μL were plated on SC-URA petri dishes and incubated for ∼48 h at 30°C to measure the transformation efficiency, aiming at 20X for each transformation (See Supplementary Table 3 for variant coverages).

The transformed pool of cells was grown for at least 24 hours before initiating the second round of plasmid selection. Input cultures were established by inoculating fresh SC–URA/MET/ADE medium at an OD600 of 0.2 from the saturated culture and incubated at 30 °C overnight. After input plasmid selection, the competition cycle started by inoculating 1L SC-URA/MET/ADE + 200 μg/ml methotrexate at an OD600 of 0.05 from the input culture. The volume equivalent to 1L at an OD600 of 1.6 from the input cultures were splitted in two equal tubes and centrifuged for 10 min at 5,000g at 4°C. Collected cells were washed twice, pelleted and stored at -20°C for later DNA extraction.

Competition cell cultures were incubated at 30 °C until reaching an OD600 of 1.6 (output), thus ∼5 generations since inoculation (see Supplementary Table 3 for OD600 values, and timings of each experiment). After incubation, output cell cultures were splitted in two equal tubes and centrifuged for 10 min at 5,000g at 4°C. Collected cells were washed twice, pelleted and stored at -20°C for later DNA extraction.

### DNA extraction, plasmid quantification and library preparation

DNA was extracted from pelleted yeast input and output stored cells using a big scale yeast DNA extraction protocol as described in^19^, scaled proportionally to the volume of harvested cells from the competition cultures. Since the final DNA samples contain both yeast genomic DNA and the target plasmids carrying the mutagenised libraries, plasmid concentrations were quantified in triplicate by qPCR. Quantification was performed using primers oGJJ152 and oGJJ153, which amplify a region within the origin of replication of the standard doubledeepPCA plasmids used in this study.

To prepare the libraries for illumina sequencing, we performed two PCR amplification cycles. In the first PCR (PCR1) we amplify the mutant library from the extracted samples while adding the illumina adapters. Primers for this were designed to add increased nucleotide complexity to the products by introducing frame-shift bases. PCRs for each library and block were performed independently with different sets of primers for each library’s constant regions, as shown in Supplementary Table 1. Primers annealing to constant regions outside of the PDZ sequences were common for each block of libraries; common reverse primer oGJJ052-58 for the block1 libraries, and common forward primer oGJJ084-89 for the block2 libraries.

In order to avoid PCR biases, each reaction was started with a minimum amount of 20X more initial template plasmid molecules than the number of illumina reads aimed per sample. These calculations accounted for the fact that multiple selection conditions were pooled within each sample. For each PCR reaction, the amount of input plasmid was estimated based solely on the representation of the specific library being amplified. For example, if a sample contained four pooled libraries assumed to be equally represented, a 4x plasmid input was used to ensure 1x coverage of the target library. Up to 50-60e6 target plasmid molecules were used per 50μL reaction of PCR1, so the number of PCR1 tube reactions was scaled according to the complexity of each library. For each sample, 15 PCR1 cycles were performed using Q5 Hot Start High-Fidelity DNA Polymerase (NEB). Excess of primers were cleaned by using 2μL of ExoSap per 50μL reaction and incubating 30 min at 37°C and 20 min at 80°C for heat inactivation. The multiple PCR1 product tubes of each sample were pooled and purified using a MinElute PCR purification kit (QIAGEN).

PCR1 product was then run through a second round of PCR (PCR2) to add the remainder of the Illumina adapter and demultiplexing indexes. Each 50 μL PCR2 reaction included 1.25 μL of PCR1 product. A total of half the number of PCR1 reactions was used in PCR2. PCR2 was run for 10 cycles at 62°C of annealing temperature using common P5 and P7 Illumina adapters with different barcode indexes. PCR products were run on a 2% agarose gel and purified using the QIAEX II Gel Extraction Kit. The final purified product was subjected to illumina paired end 2x150 sequencing. For all downstream sequencing analyses, we targeted a minimum median coverage of 100 reads per variant

Alongside the preparation of all mutagenised libraries, the same protocol was applied to the original wild-type plasmids. These were sequenced in parallel to enable later correction of variant counts based on wild-type sequencing data.

### Sequencing data processing

FastQ files from all libraries and all blocks were processed using DiMSum pipeline^46^. FASTQ files corresponding to the same sample but originating from different sequencing runs or lanes were treated as technical replicates. Two rounds of DiMSum runs were performed. In the first round, all input and output samples were analyzed together with wild-type sequencing data using default DiMSum parameters.

Following this initial run, variant counts were corrected based on the wild-type sequencing. Variants observed in the wild-type plasmid, primarily single-nucleotide substitutions, are assumed to arise from sequencing or PCR errors. Their calculated proportions with respect to the wild type were removed from the datasets, under the assumption that they result from sequencing errors of the abundant wild-type background.

Once the new variant count files were calculated, the next dimsum run was performed. Restarting at stage 4, the modified variant count files were given as input. Together with this, barcode-variant identity files were also provided to restrict the analysis to designed variants only (option “barcodeIdentityPath”). In the cases where the nucleotide wild-type variant was not present, the most abundant synonymous variant was selected as the new wild type. Additional input count filters (“fitnessMinInputCountAny”) were manually adjusted for each library based on the first DiMSum run’s report of variant count distributions, aiming to minimise the proportion of reads per variant attributed to sequencing errors of lower order mutants. Filtering thresholds were informed by the input count distribution plots (Stage 4.1) and expected read counts of variants arising from sequencing errors, included in the DiMSum report^46^.

### Normalization of fitness scores across experiments

Normalised and processed fitness scores were only used for visualization purposes in Fig. 1 and Extended Data Fig. 1. To ensure comparable fitness value ranges, those were first normalised between blocks of the same protein library, and further normalised across different assays.

Normalization was performed by using stop codon variants, which represent the lowest expected fitness values, and synonymous variants, expected to have fitness around 0, as lower and upper bounds, respectively. For each protein, we calculated the median fitness of all stop codon variants in the overlapping region between blocks (regardless of background mutations), along with the median of all synonymous variants. If synonymous variants were not present, the upper bound was set to zero. A linear regression model was then used to scale the fitness values of each block relative to block 1, which was taken as the reference. After normalizing within blocks, a second linear regression model was applied to scale fitness effect values across all assays. The PSD95-PDZ2 binding NMDAR2A assay was used as the reference. This model was based on the median fitness values of all stop codon variants and all synonymous variants from both blocks.

### Free energy inference with MoCHI

We used MoCHI v0.9^20^ to fit global mechanistic models to the complete dataset, including 22 doubledeepPCA assay datasets accross multiple PDZ domain homologs and their respective binding partners.

We fitted five models, one per PDZ domain. Each model was fitted by simultaneously incorporating selection data from both AbundancePCA and BindingPCA assays and both mutagenised blocks. For domains with a single binding partner, four datasets were included in the fit, two AbundancePCA blocks and two BindingPCA blocks. For domains with two binding partners, six datasets were fitted simultaneously, consisting of two abundancePCA blocks and two bindingPCA blocks for each interaction. This model assumes that folding free energy changes are independent of the bound ligand, allowing folding energies to be shared across interactions. Consequently, for PDZ domains with multiple binding partners (PSD95-PDZ2 and NHERF3-PDZ1), a single folding energy model was fitted simultaneously with separate binding datasets for each interaction.

In this model, the protein is assumed to exist in three distinct states: unfolded and unbound, folded and unbound, and folded and bound. The probability of the unfolded and bound state is considered negligible. This model fits single mutation Gibbs free energy of folding (ΔG_f_) and binding (ΔG_b_) effects, and works under the assumption that the changes in free energy of variants (ΔΔG_f_ and ΔΔG_b_) are additive. This means that the effect of mutations from a multi-mutant are simply the sum of effects of the single mutations.

The mochi parameters used were set to default as explained in previous work^19,20,22^. Each model architecture consisted of one additive layer representing folding energy, and one additive layer per binding assay. Accordingly, models for PSD95-PDZ3, NHERF3-PDZ2, and ERBIN-PDZ1 included two additive layers (one for folding and one for binding), while models for PSD95-PDZ2 and NHERF3-PDZ1, which have two binding partners, included three additive layers. The non-linear transformations used, “TwoStateFractionFolded” (for AbundancePCA data) and “ThreeStateFractionFolded” (for BindingPCA data), were derived from the Boltzmann distribution, relating fraction of protein folded/bound to the underlying free energy changes.

The input to the models consisted of raw fitness scores from both single and double mutants, calculated prior to any normalization. Each dataset was manually adjusted so that variant entries contained the full-length amino acid sequence of the corresponding PDZ domain, rather than just the mutated region of the block.

### Protein structures, structure metrics, and contacts

The PDZ domain structures, alone and in complex with the ligands, were all obtained from AlphaFold3 predictions^44^. Given the low plDDT score of the predicted C-terminal tail extension from NHERF3-PDZ2 (70 > plDDT > 50), for visualization purposes it was manually adjusted to remove the secondary structure annotation, reflecting as well prior evidence of its intrinsic disorder and dynamic nature in NMR PDB entry for the solution structure of Second PDZ domain of PDZ Domain Containing Protein 1 (PDB ID: 2EEI; Niraula et al., 2008, unpublished)^47^.

Relative solvent accessible surface area was calculated for each unbound structure using freeSASA python package (v2.2.1)^48^. All residues with rSASA<0.25 were defined as core.

Minimum ligand-residue distances were calculated as the shortest distance between any side chain heavy atom of the respective PDZ residue and any ligand side chain heavy atom. Residue contacts were defined using getContacts (https://getcontacts.github.io/) on the AlphaFold3 predicted structures, by using get_static_contacts.py with parameters --itypes all.

Finally, binding interface residues were defined as those with at least one predicted contact with the ligand.

### Structure visualizations

ChimeraX^49^ was used to visualise all protein structures presented. Contacts in these structures display the predictions from getContacts(https://getcontacts.github.io/), which are drawn using the distance command.

We used SSDraw^50^ for the 2D secondary structure representation of the mutated PDZ domains, using as input the corresponding AlphaFold3 structure predictions.

### Structural alignments

Structural alignments were performed using T-Coffee^51^ Expresso with default parameters. Residue conservation was calculated from the structural alignment using the BLOSUM62 substitution matrix to quantify residue similarity.

### Classification of binding interface hotspots

Hotspots were defined as interface residues whose distribution of ΔΔG_b_ values was significantly greater than the median ΔΔG_b_ value across all binding interface residues (0.75kcal/mol). Statistical significance was assessed using a one-sided t-test, and p-values were corrected for multiple testing using the false discovery rate (FDR) method Benjamini-Hochberg, with a significance threshold of *FDR*<0.1. Classification of hotspots can be found in supplementary table 6.

### Quantification of distance dependence effects of mutations

To quantify the dependence of mutation effects on the distance from the ligand, we computed the minimum heavy atom side chain distance between each residue and the ligand, using AlphaFold3-predicted^44^ structures. To quantify this dependence, an exponential decay function was fitted to the data using the nls() function from the R stats package, with initial parameter estimates obtained via the optim() function. In exponential decay model *y = a · e^bx^*, *a* represents the estimated |ΔΔG_b_| at distance 0 Å from the ligand, *b* is the decay rate and x is the minimum heavy atom side chain distance to the ligand. The model was fit to all data points excluding those corresponding to binding interface residues.

The same analysis procedure was applied to previously published datasets for KRAS^22^, GB1 and GRB2^19^, using previously defined annotations. For these datasets, distances were calculated as the minimum side chain heavy atom distance to each interaction partner, and binding interfaces were defined as any residue within 5 Å of the interaction partner.

Similarly, the procedure was applied to SRC data^23^, where distances were annotated as the minimum distance between each residue’s Cα and three functional features: (1) the non-hydrolyzable ATP analog AMP-PNP (from PDB: 2SRC), (2) the catalytic residue D388, and (3) the proposed phosphosite substrate positioning residue P428. Binding interface residues for SRC were defined based on previously annotated active sites from the literature^23^.

### Classification of allosteric hotspots

To identify allosteric hotspots, we first calculated the expected ΔΔG_b_ at each residue based on its minimum side-chain heavy atom distance to the ligand, using the previously fitted exponential decay model. Residuals were then computed as the difference between observed and expected ΔΔG_b_ values. Residues were classified as allosteric hotspots if their residual distribution was significantly greater than zero, using a one-sided t-test with FDR correction (*FDR* < 0.05). To avoid false positives from small effects at long distances, we further required a median of the absolute ΔΔG_b_> 0.2. Classification of hotspots can be found in Supplementary table 6.

## Acknowledgements

This work was funded by Wellcome (220540/Z/20/A), European Research Council (ERC, Advanced Grant 883742), Spanish Ministry of Science and Innovation (LCF/PR/HR21/52410004, EMBL Partnership, Severo Ochoa Centre of Excellence), AGAUR (2021 SGR 01226), and the CERCA Program/Generalitat de Catalunya. A.M.-A. was funded in part by a fellowship from ”laCaixa” Foundation (ID 100010434, fellowship code B006052). We thank all members of the Lehner Lab for helpful discussions and suggestions.

## Data availability

All DNA sequencing data have been deposited in the European Nucleotide Archive (ENA) at EMBL-EBI under accession number PRJEB90582

## Code availability

Source code to reproduce the analyses is available at https://github.com/lehner-lab/PDZ_homologs

## Author contributions

A.M. performed all experiments and analyses. A.M. and B.L. designed experiments and analyses and wrote the manuscript.

## Competing interests

B.L. is a founder and shareholder of ALLOX.

---

**Extended Data Figure 1: Energetic landscapes of multiple PDZ domains homologs**

**a.** Triplicate fitness Pearson correlations and fitness density distributions for all libraries, including AbundancePCA, BindingPCA, and two blocks each. **b.** Pearson correlation of fitness measurements in overlapping regions between blocks (i.e. repeated variants measurements in independent experiments). Fitness values are shown after normalization between blocks and libraries. Red scatter points indicate libraries lacking single amino acid substitutions for the wild-type background; for visualization, data from the most wild-type-like background is shown (ΔΔG_b_ of the background ∼0). Error bars indicate 95% CI (n = 3 biological replicates). **c.** Fitness heat maps for single mutants for AbundancePCA and BindingPCA results. As in panel b, for blocks lacking single variants on the wild-type background, data is shown for the most wild-type-like background. Dashes indicate wild-type residues. Residue numbering follows UniProt protein sequence annotation **d.** Scatter plots comparing abundancePCA and bindingPCA fitness measurements for single amino acid substitutions from the same PDZ domain.

**Extended Data Figure 2: From molecular phenotypes to free energy changes**

**a.** Performance of models fit to ddPCA data for all libraries. Pearson correlation is reported per library and block. **b.** Non-linear relationships (global epistasis) between observed AbundancePCA fitness and changes in free energy of folding. **c.** Non-linear relationships (global epistasis) between observed BindingPCA fitness and both changes in free energy of folding and binding **d.** Free energy changes distributions of core vs surface residues. “****” annotation refers to *P*<2.2x10-16 for Wilcoxon rank-sum test. **e.** Scatter plots comparing folding and binding free energy changes for single amino acid substitutions for the same PDZ domain.

**Extended Data Figure 3: Binding interface energy landscapes for seven interactions**

**a.** Structure representation of hotspots in every interaction. All highlighted residues are ligand contacts. Colors indicate non-hotspots (white), hotspots in 1 PDZ domain (purple), and recurrent hotspots in at least 3 (red). Ligand is represented in light grey. Dashed black lines show all predicted structural contacts with the ligand. **b.** Heat map of all structurally aligned residues that are classified as binding interface hotspots in at least one interaction. Letter labels refer to wild type residues in each position and interaction. Scatter points overlaid on the heat map represent ligand wild-type amino acids that make direct contact with the corresponding PDZ domain residue. Scatter points below the heat map represent the residue’s minimum side chain heavy atom distance to the ligand. **C.** Same heat map for residues never classified as binding interface hotspots.

**Extended Data Figure 4: Seven comparative allosteric maps**

**a.** Relationships between the absolute ΔΔG_a_ and the minimum distances to the binding partner. Lines represent exponential decay fits (*y = a · e^bx^*), calculated excluding mutations within the binding interface for SRC, **b.** for KRAS and all interaction partners, and **c.** for GB1 and GRB2-SH3 domains. **d.** Pearson correlation of ΔΔG_b_ residuals across all pairwise comparisons of libraries. **e.** Percentage of shared mutations with significantly positive ΔΔG_b_ residuals (*FDR*<0.1, one-sided Z-test) for each pairwise library comparison. **f**. Boxplots of ΔΔG_b_ values for all residues across all tested interactions. For each residue, the blue dot indicates the expected ΔΔG_b_ based on its distance to the ligand, derived from interaction-specific distance decay fits. Boxplots outlined in black represent conserved allosteric hotspots found in three or more PDZ domains (*FDR* < 0.05, one-sided t-test); green outlines indicate hotspots present in one or two PDZ domains. Red box plots correspond to binding interface residues. **G**. Distribution of ΔΔG_b_ values per interaction across five residue categories: non-hotspots outside the binding interface (gray), allosteric hotspots found in one or two PDZ domains (green), conserved allosteric hotspots found in three or more (purple), binding interface hotspots (orange), and binding interface non-hotspots (yellow). “*” indicate *P*<0.05 (Wilcoxon rank-sum test) **h**. Enrichment test for identified hotspots defined by previously reported allosteric networks in PSD95-PDZ3. Bars represent the most significant result out of four one-sided Fisher’s exact tests conducted per dataset. Tests include: enrichment of hotspots identified in PSD95-PDZ3 binding CRIPT, excluding this interaction’s binding interface residues (blue); enrichment of conserved hotspots (i.e., those identified in at least three PDZ domains in this study), excluding PSD95-PDZ3 binding interface residues (green); enrichment of hotspots identified in PSD95-PDZ3 binding CRIPT, including binding interface residues and hotspots (orange); and enrichment of the most conserved hotspots, including binding interface residues (purple). **i.** One-sided Fisher’s exact test enrichments of allosteric hotspots in core (rSASA<0.25) and surfaces, calculated separately for each interaction. **J**. ΔΔG_b_ Pearson correlation for PSD95-PDZ2 binding two different partners; error bars indicate 95% confidence intervals from a Monte Carlo simulation approach (n = 10 experiments), and the same for **k**. NHERF3-PDZ1. **L.** ΔΔG_b_ Pearson’s correlation for ΔΔG_b_ of mutations in hotspots present in one or two PDZs.

**Extended Data Figure 5: Gain of function allosteric sites.**

**a.** Relationship between absolute ΔΔG_b_ and the minimum side chain heavy atom distance to the ligand for gain-of-function (ΔΔG_b_<0, top) and loss-of-function (ΔΔG_b_>0, bottom) mutations. Curves show exponential decay fits (*y = a · e^bx^*), calculated excluding mutations within the binding interface. **b**. Number of mutations per residue with ΔΔG_b_< -0.2 and significantly ΔΔG_b_<0 effects (FDR<0.1, one-sided Z-test). Blue bars indicate surface residues; grey bars indicate core residues (rSASA<0.25). Black outlines mark residues significantly enriched in GOF mutations (FDR<0.1, one-sided FET). **c**. Structural visualization of residues enriched in mutations with significantly ΔΔG_b_<0 (FDR<0.1, One-sided FET).

